# Estimating chromosome sizes from karyotype images enables validation of *de novo* assemblies

**DOI:** 10.1101/2022.05.22.492982

**Authors:** Arne Ludwig, Alexandr Dibrov, Gene Myers, Martin Pippel

**Author notes:** Equal contribution (Arne Ludwig, Alexandr Dibrov). To whom correspondence should be addressed: Martin Pippel, Max Planck Institute of Molecular Cell Biology and Genetics, Pfotenhauerstr. 108, 01307 Dresden, Germany, Mail.

## Abstract

Highly contiguous genome assemblies are essential for genomic research. Chromosome-scale assembly is feasible with the modern sequencing techniques in principle, but in practice, scaffolding errors frequently occur, leading to incorrect number and sizes of chromosomes. Relating the observed chromosome sizes from karyotype images to the generated assembly scaffolds offers a method for detecting these errors.

Here, we present KICS, a semi-automated approach for estimating relative chromosome sizes from karyotype images and their subsequent comparison to the corresponding assembly scaffolds. The method relies on threshold-based image segmentation and uses the computed areas of the chromosome-related connected components as a proxy for the actual chromosome size. We demonstrate the validity and practicality of our approach by applying it to karyotype images of humans and various amphibians, birds, fish, insects, mammals, and plants. We found a strong linear relationship between pixel counts and the DNA content of chromosomes. Averaging estimates from eight human karyotype images, KICS predicts most of the chromosome sizes within an error margin of just 6 Mb.

Our method provides additional means of validating genome assemblies at low costs. An interactive implementation of KICS is available at https://github.com/mpicbg-csbd/napari-kics.

## Introduction

Highly contiguous, complete, and accurate genome assemblies are fundamental to associating genotypes with phenotypes [19, 22]; genome-based evolution [19] and speciation studies [22]; analyzing repeat-organization and function [28]; population genetics [33]; and, ultimately, biomedical research [29, 47]. *De novo* assemblies can achieve chromosome-length scaffolds using third-generation long sequencing reads combined with additional sequencing data for scaffolding such as Bionano optical maps or Hi-C chromatin interaction maps [24, 40, 31].

However, the resulting assemblies often contain scaffolding errors that must be manually curated [18]. Knowing the true chromosome sizes in this step helps identify severe misjoins and incomplete scaffolds. An open-source tool to estimate chromosome sizes would assist in the *de novo* genome assemblies of unsequenced species like those targeted by the Vertebrate Genome Project [8], the Bird 10 000 Genomes Project [48], and the Earth BioGenome Project 2020 [27].

Karyotyping is a well-established technique in cytogenetics [13] using photomicrographs of complete chromosome sets. It has been practiced for more than a century [13] producing karyotype images for hundreds of species. Such data distinctly renders the individual chromosomes’ outlines and can thus be used to estimate their morphological properties. Commercial software packages like LUCIA Karyo [26] or Ikaros Karyotyping Platform [30] offer karyotype segmentation and subsequent analysis capabilities. We would argue that the scientific field would benefit from an open-source tool providing similar features.

Here, we present the karyotype image-based chromosome size estimator (KICS), a semi-automated method to estimate relative chromosome sizes from karyotype images. The open-source tool is implemented as a plugin for the general-purpose image viewer napari [9] and is available at https://github.com/mpicbg-csbd/napari-kics.

## Results

### Method Overview

The semi-automated method presented in this paper estimates relative chromosome sizes from karyotype images in four steps: (1) initial image segmentation by thresholding, (2) labeling of connected components, (3) manual curation of image labels, and (4) (semi-)automatic naming, ordering, and grouping of chromosomes. Optionally, the user may provide an estimate for the haploid genome size in order to derive absolute chromosome sizes. The results are available as an annotated image and in tabular format for further analysis. Figure 1 gives an overview of the process. We implemented the workflow as an extension to the general-purpose image viewer napari [9], which supplies functionality to load and annotate images.

**Figure 1:**
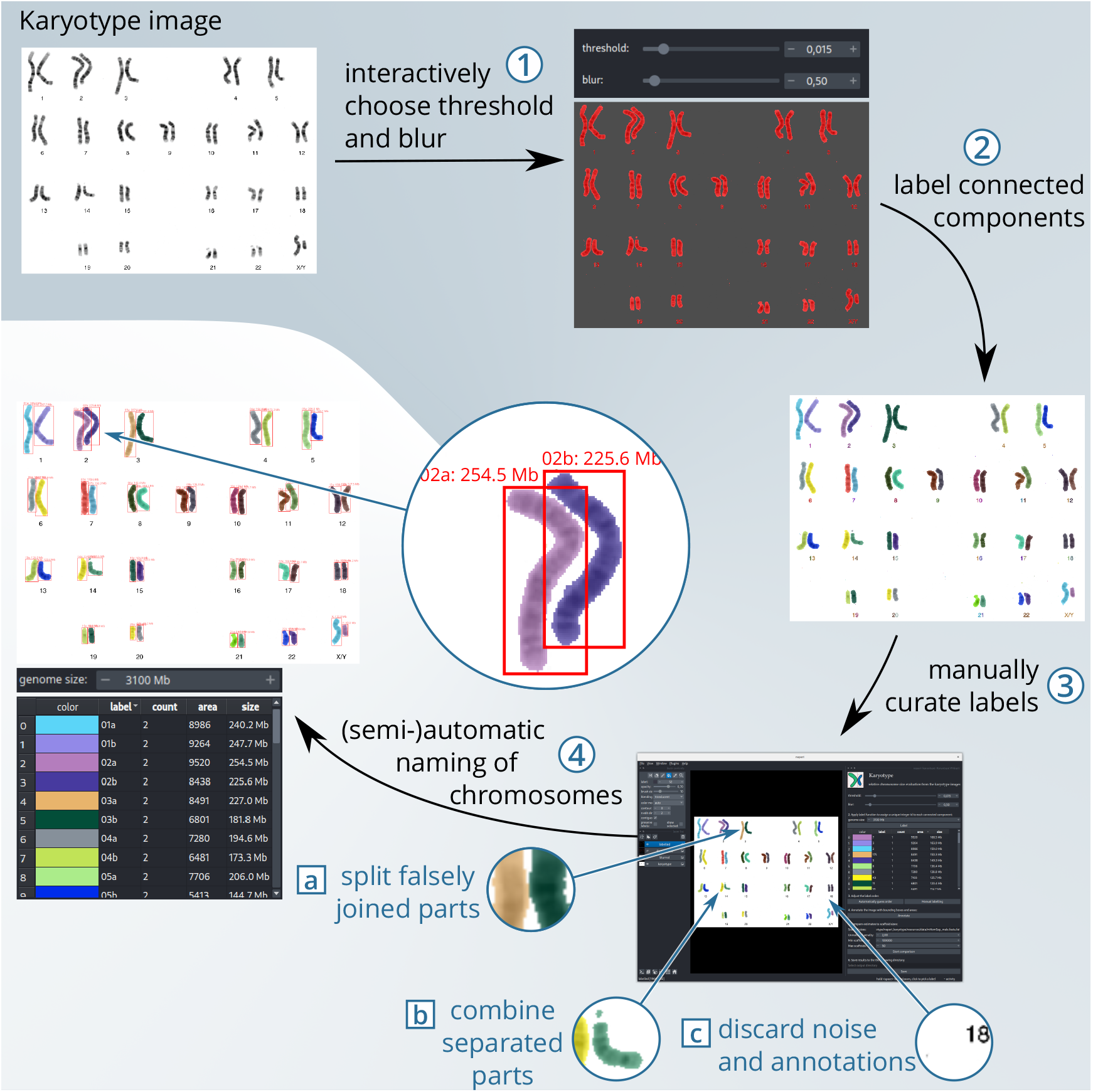
Overview of the workflow for estimating chromosome sizes from a karyotype image.

### Quality Analysis

To evaluate the accuracy and precision of KICS, we tested it on a set of eight human karyotypes, Human8, which provides consistently high-quality images of 367 chromosomes and accurate reference chromosome sizes from the novel human telomere-to-telomere assembly [37]. We generated the estimates with a threshold of *θ* = 0.05, blurring radius *σ*_B_ = 1, and genome size *G* = 3.1 Gb. In manual curation, we only joined falsely separated pieces of chromosomes (fig. 1b) and removed noise-induced objects (fig. 1c). We named the chromosomes using the automatic method and re-named the sex chromosomes to X and Y, as indicated in the images. The dataset includes eight chromosomes with major translocations and deletions. These are ex-cluded from statistical analyses but highlighted in the plots demonstrating the potential of our method to reveal unexpected deviations. The resulting estimates are presented in figure 2.

**Figure 2:**
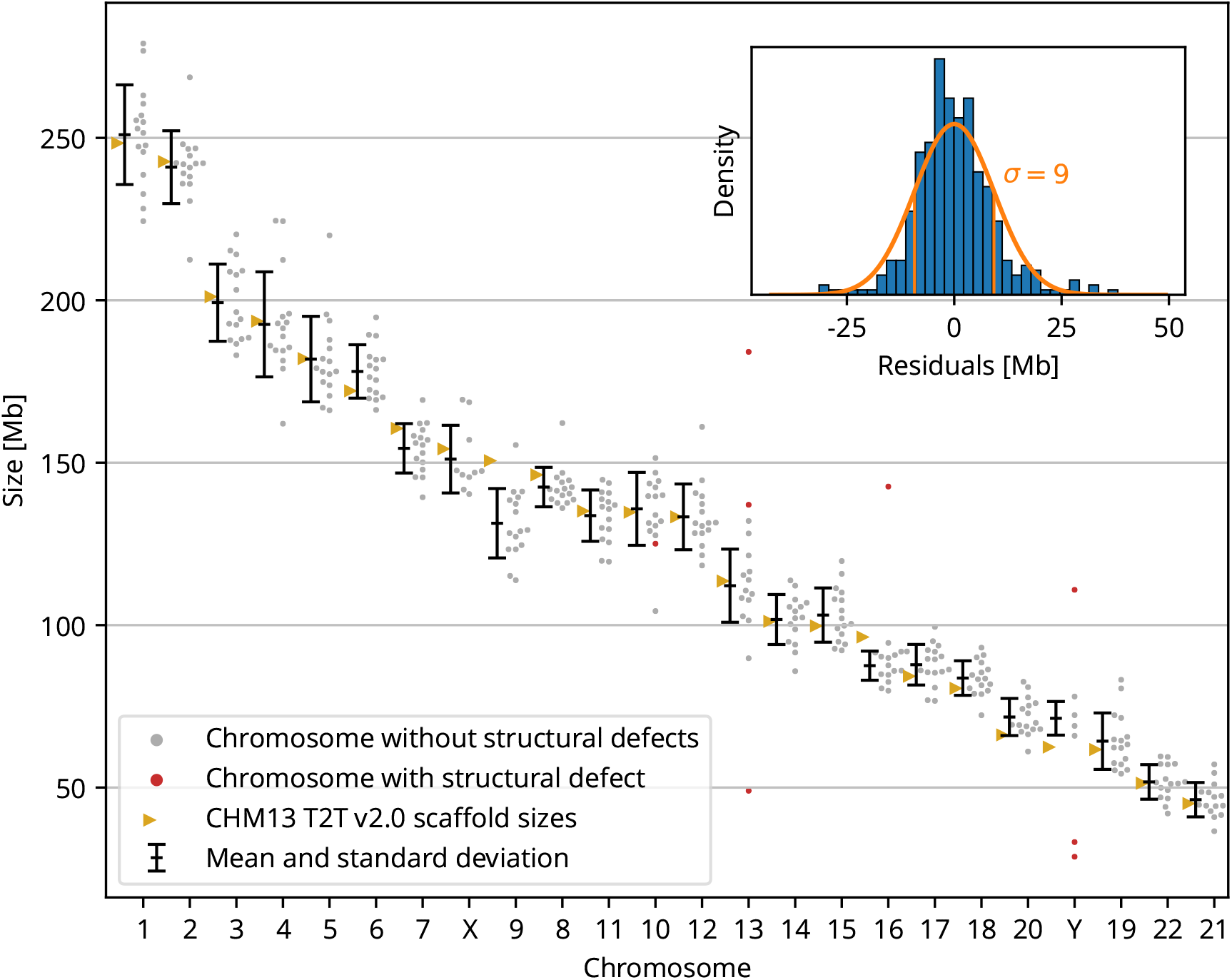
Estimated sizes of individual chromosomes with mean and standard deviation. Inset: distribution of residuals (histogram) and the inferred normal distribution (orange curve). Chromosomes with structural defects (red dots) are not included in the evaluation but highlight the potential of KICS to reveal unexpected deviations.

In the following, we examine the accuracy and precision of KICS. While *accuracy* describes systematic errors in the estimates, *precision* is a measure of their statistical spread. We define the *accuracy* of the estimates as the absolute difference between their mean and the reference value; and their *precision* as their standard deviation.

KICS estimates chromosome sizes to a high degree of accuracy, mostly within 6 Mb of the reference size. On a per-chromosome basis, the method achieves an accuracy of 4.2%or better relative to the true chromosome size. Exceptions of the above accuracy are chromosomes 9 (−19 Mb/−13%), 16 (−9 Mb/−9%), 20 (−6 Mb/−8%), and Y (+9 Mb/+14%). The variations of these estimates could be due to the satellite sequences in the centromeres [3]. Satellite sequence gets tightly packed and may therefore appear smaller in karyotype images. For example, chromosomes 9 and 16 are underestimated by KICS and contain about 20%and 16%of satellite sequence, respectively [3]. On the other hand, the length of chromosome Y is overestimated which might indicate missing sequence. Despite these exceptions, in general KICS is still able to estimate chromosome sizes from karyotype images with a high degree of accuracy.

To analyze the precision of the estimates, we consider the distribution of residuals across all estimates. The resulting histogram visually agrees with a normal distribution (fig. 2). Thus, for practical purposes, we derive a normal-distributed error with mean *μ* = 0 and standard deviation *σ* = 9Mb. Considering the precision of individual chromosomes, we found a pronounced correlation between chromosome size and precision (supp. fig. 2C), indicating a multiplicative error model that we discuss in depth below.

Because chromosome condensation generally differs between species [23, 10, 5], our method may not be equally valid for all organisms. Therefore, we evaluated it on a wide range of species covering amphibians *(A. mexicanum, X. laevis*), birds (G. *gallus*), insects *(D. melanogaster),* plants (A. *thaliana, Z. mays*), fish *(D. rerio*), and mammals (*R. norvegicus*, *H. sapiens*), by correlating estimated with reference chromosome sizes (fig. 3). For individual species, we found Pearson correlations from slightly significant (A. *thaliana*, *ρ* = 0.77) to highly significant (A. *mexicanum*, *D. melanogaster*, *G. gallus*, *H. sapiens*, *R. norvegicus*; 0.97 ≤ *ρ* ≤ 0.98). We observed that the correlation coefficient *ρ* is mainly influenced by the quality of the karyotype images, which we go into more deeply in the discussion. Overall, the estimates and reference sizes agree across species.

**Figure 3:**
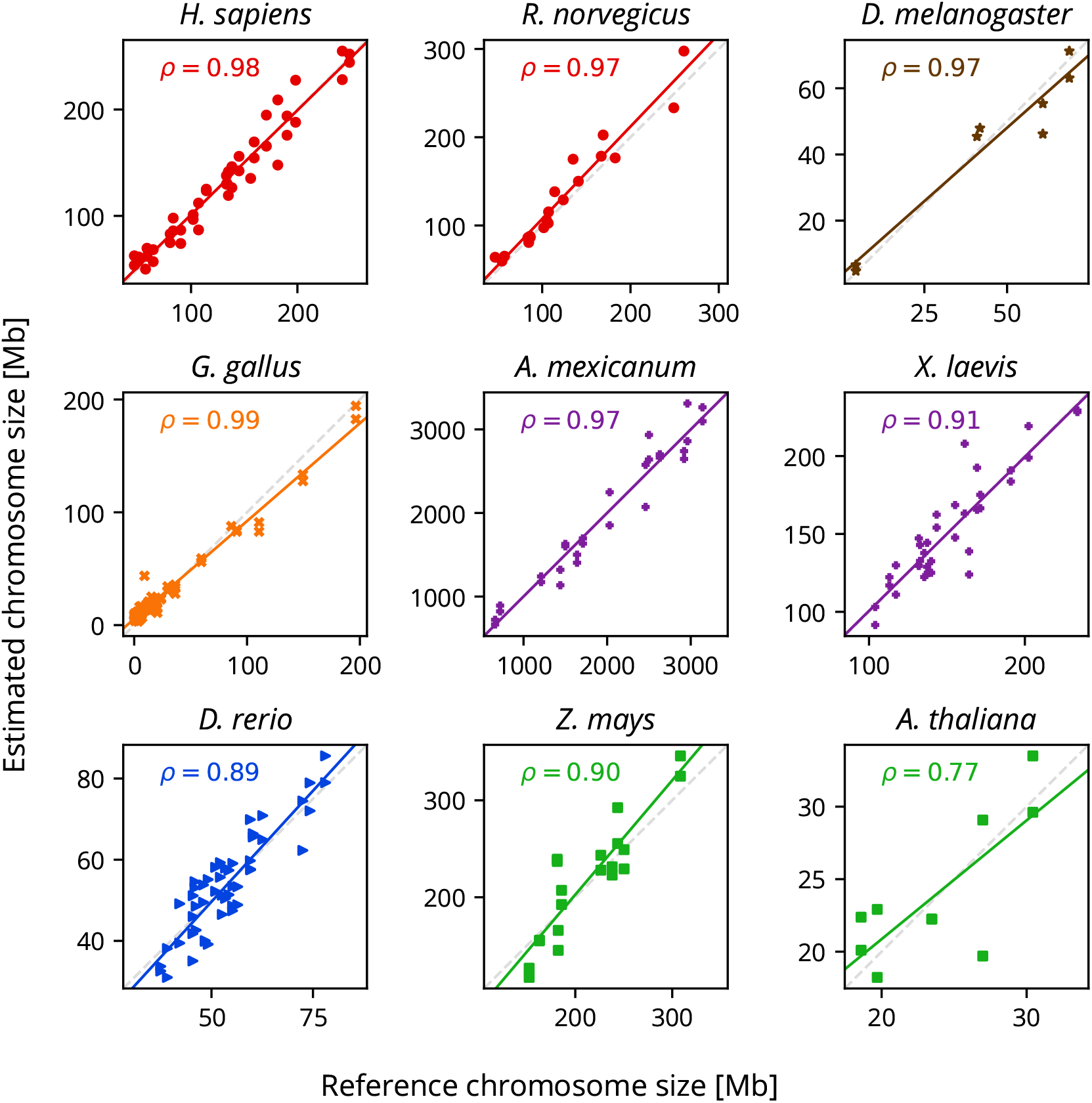
Reference versus estimated chromosome size for mammals (red circles), insects (brown stars), birds (orange crosses), amphibians (purple pluses), fish (blue triangles), and plants (green squares). Linear regression is depicted by a solid line and the identity function by a dashed line for reference. *ρ* shows the Pearson correlation coefficient.

### Influence of Threshold

The thresholding value *θ* determines the segmentation and, consequently, may distort the chromosome size estimates. Therefore, we investigated the ease of choosing a near-optimal value and its robustness against perturbations. We used the Pearson correlation between the estimated and reference chromosome sizes as a measure of fit and evaluated it for different values of 0 ≤ θ < 1 and *σ*_B_ = 1 on the Human8 dataset (fig. 4A). These results suggest an optimal threshold around *θ*\* = 0.02 for all karyotypes and that any value 0.01 ≤ *θ* ≤0.20 (fig. 4) produces similarly good results. Without prior knowledge of this analysis, the manual choice *θ* = 0.05 was close to the designated optimum *θ*\*. However, these results also suggest that the threshold is volatile towards zero. This agrees with the visual effects of low thresholds (fig. 4B), which the user easily detects. On the high end of the near-optimal interval, a threshold of *θ* = 0.20 yields drastically impaired segmentation results and would be discarded by the user. Overall, the choice of the threshold is straightforward and robust.

**Figure 4:**
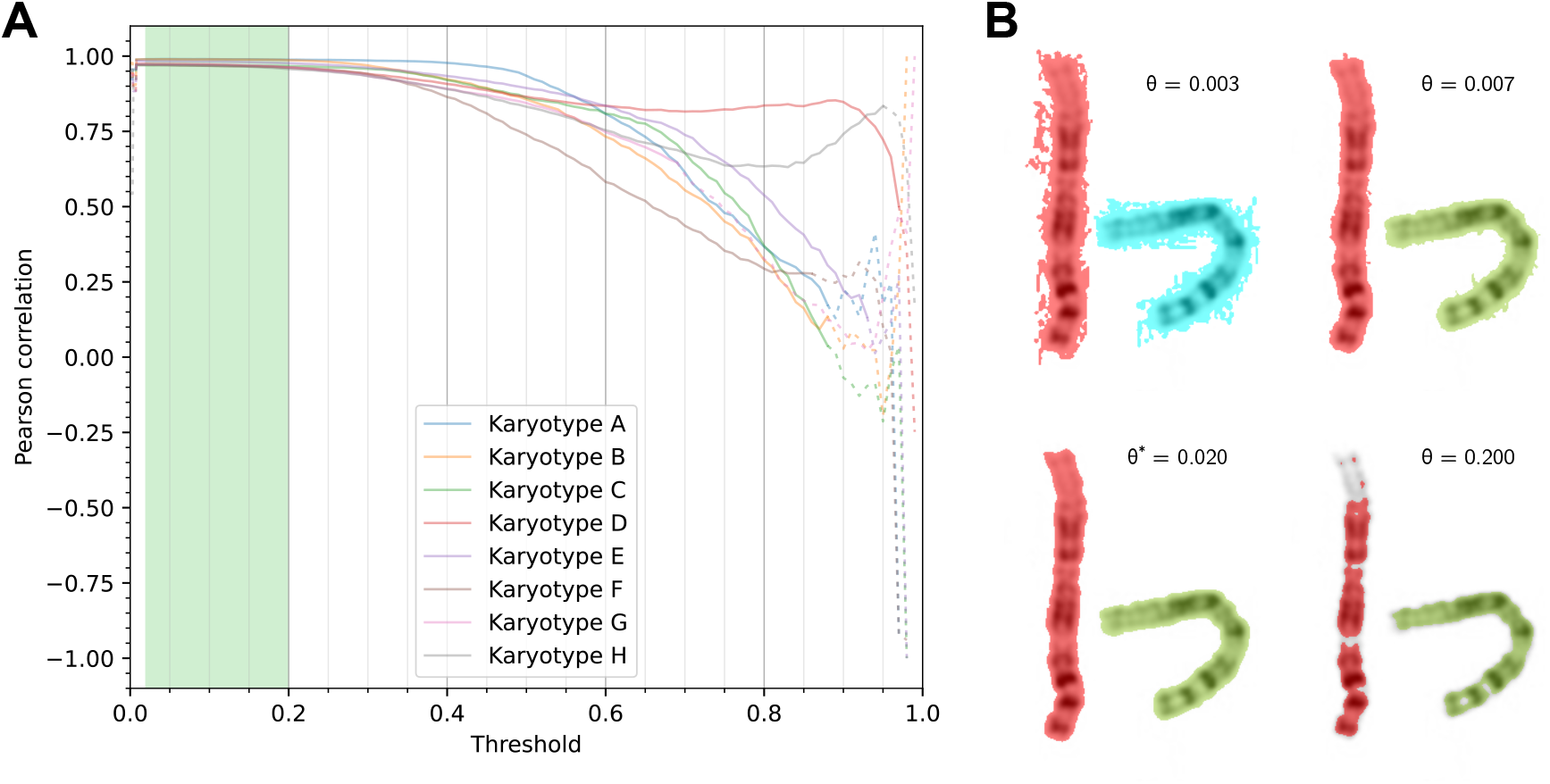
**A)** Pearson correlation of estimated and reference chromosome sizes at varying values of the threshold parameter *θ* for the Human8 dataset. Dashed lines show segmentation into incorrect numbers of chromosomes. Highlighted in green is the near-optimal parameter space. **B)** Segmentation results of chromosome pair 1, karyotype H at varying thresholds *θ*.

### Examples

We wished to test KICS in the context of de novo genome assembly, where chromosome sizes are unknown *ab initio*. We calculated the estimates from single karyotype images and compared with manually curated chromosome-scale assemblies for four species – one fish, one bat and two birds. Because the true matching is unknown, we order the estimates and scaffold sizes independently by size from largest to smallest and match the numbers by rank.

The first example is the recent assembly of the West African lungfish *(Protopterus annectens*) which has an exceptionally large genome size of 40.5 Gb [46]. As can be seen in figure 5A, both number series have similar characteristics: (1) the two largest sizes are distinctly larger than the rest, (2) the three smallest sizes are very similar, and (3) the remaining twelve sizes have a roughly linear slope. Because there is no severe discrepancy between the number series, we cannot invalidate the scaffold sizes. In other words, the chromosome size estimates support the scaffold sizes.

**Figure 5:**
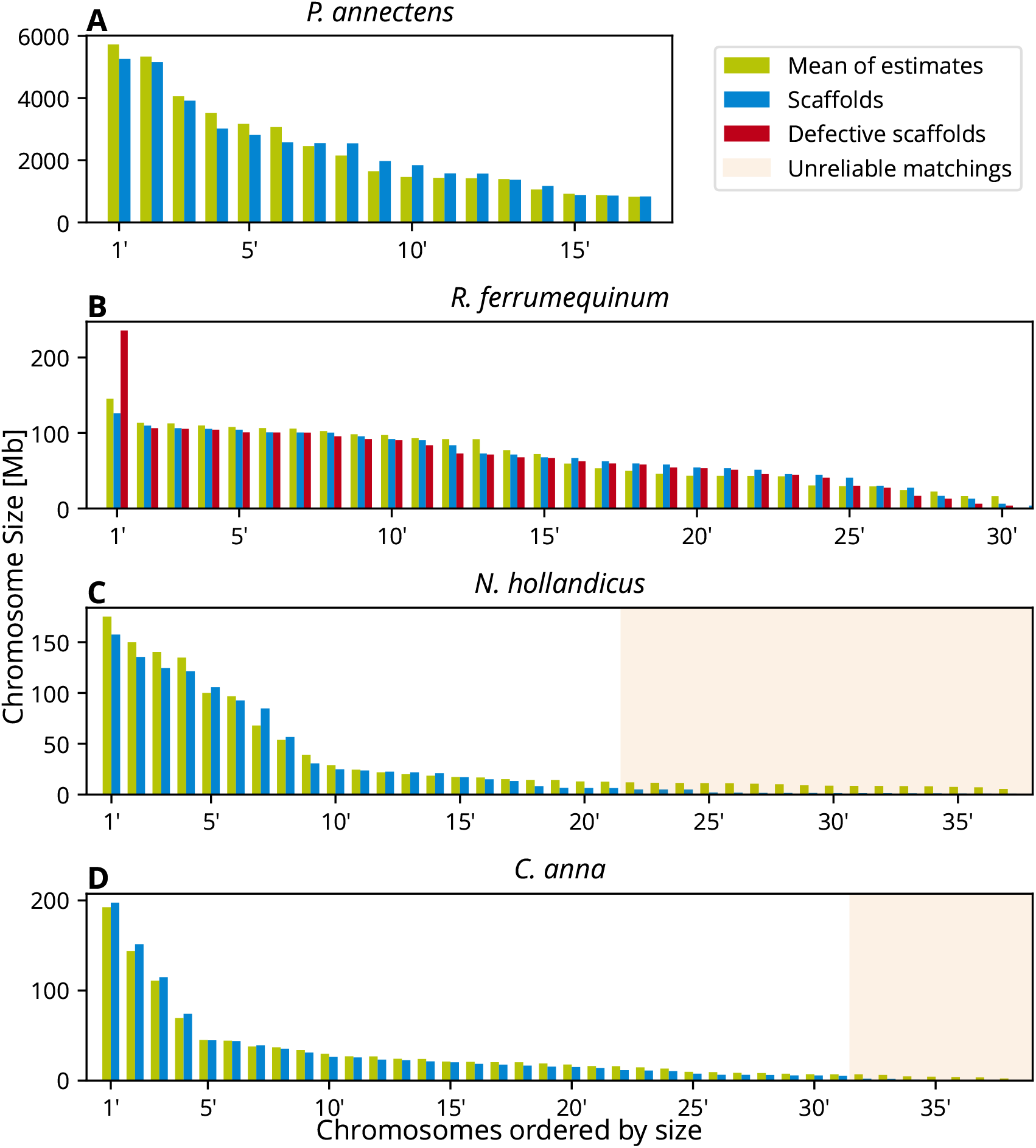
Comparison of chromosome size estimates and de novo assembly scaffold sizes without knowledge of correct matching. Both estimates and scaffold sizes are sorted independently by size before plotting. The hatched area highlights matches that are deemed unreliable, i.e. where the scaffolds have less than half the size of the corresponding estimate.

In our second example, we show a defect that we encountered assembling the genome of the greater horseshoe bat *(Rhinolophus ferrumequinum)* published in [19]: two of the largest chromosomes were falsely joined at the telomeres by the Hi-C scaffolding procedure. In figure 5B, the defective scaffold sizes are shown in red, while the corrected sizes are shown in green. The size of the falsely joined scaffold distinctly stands out because it has roughly double the size of the largest chromosome estimate. After manual curation of the scaffolds, both number series neatly align except for the last pair, possibly indicating missing sequence in one of the smallest chromosomes.

The last two examples (fig. 5D and C) highlight challenges associated with microchromosomes. Microchromosomes alongside high diploid numbers of 2n ≈80 chromosomes are commonly encountered in birds [12], such as the cockatiel (*Nymphicus hollandicus, 2n* = 72) and Anna’s hummingbird *(Calypte anna, 2n* = 64). The assembly of *N. hollandicus* is an internal work in progress, and about half of the microchromosomes appear to be incomplete or missing. On closer examination, the seventh chromosome estimate is noticeably larger than the corresponding scaffold size. This may have at least two reasons which cannot be distinguished without further data: (1) a false join between one of the macrochromosomes and one of the microchromosomes, or (2) underestimated chromosome size (see human chromosomes 9 and 16, fig. 2). This illustrates an intrinsic limitation of this method: minor errors are virtually undetectable. Generally, though, the sizes of microchromosomes are estimated without notable bias, as demonstrated by the last example, *C. anna,* where all but seven microchromosomes were successfully assembled.

## Discussion

We have presented KICS, a novel semi-automated method for estimating relative chromosome sizes from a karyotype image. The method was developed with the primary goal of providing additional means of validating *de novo* genome assemblies. Our analysis based on eight karyotype images of human individuals shows that KICS accurately estimates most chromosome sizes within an error margin of just 6Mb and precision of 9Mb. Further, we demonstrated that KICS performs well on a wide variety of species by applying it to karyotypes of amphibians, birds, fish, insects, mammals, and plants. We provided evidence that the methods’ main parameter, the threshold, is straightforward to determine and robust against perturbations. Finally, we presented four practical examples demonstrating the power and limitations of KICS.

The quality of the karyotype image is the most influential variable of KICS (supp. fig. 1A). High image contrast and uniform background are vital to a good threshold-based segmentation. Low-quality images may still be used but require increased efforts for manual segmentation. Another important consideration is the copy number of the chromosomes in the karyotype image and the scaffolds; they should, e. g., contain the same set of sex chromosomes. For some species, this may be even more complex, e.g., in the freshwater planarian *(Schmidtea mediterranea)* a large translocation between two chromosomes determines whether they reproduce sexually or asexually [36]. Microchromosomes are especially hard to estimate because their size is closer to the image resolution and they easily get out of focus [20].

An alternative approach to estimating chromosome sizes from existing cytogenetic analyses is to use length estimates from the literature directly. For example, we found appropriate estimates for the chicken (*G. gallus*) in [21], table 1. We computed estimates for the chromosome size from the mean relative lengths and converted them to base pairs analogous to our method (supp. fig. 1D). We found that both methods yield similar results (supp. fig. 1C). Thus, if available, chromosome length estimates from previous studies may provide a viable alternative to our method.

**Table 1:** Listing of data sources for the Diversity dataset. Chromosome sizes were acquired from the assemblies referenced by their GenBank accession number if given. *Chromosomes were arranged according to [45, fig. 1].

| Group | Species | Karyotype image | Chromosome Sizes |
| --- | --- | --- | --- |
| Amphibians | *Ambystoma mexicanum* | [6, fig. 1]* | [45, fig. 1] |
|  | *Xenopus laevis* | [42, fig. 1c] | GCA_017654675.1 |
| Birds | *Gallus gallus* | [20, fig. 3] | GCA_016699485.1 |
| Fish | *Danio rerio* | [11, fig. 1A] | GCA_000002035.4 |
| Insects | *Drosophila melanogaster* | [17, fig. 1] | [1, fig. 1] |
| Mammals | *Homo sapiens* | [35] | GCA_000001405.28 |
|  | *Rattus norvegicus* | [16, fig. 1] | GCA_015227675.2 |
| Plants | *Arabidopsis thaliana* | [14, fig. 1] | GCA_000001735.2 |
|  | *Zea mays* | [41, fig. 1A] | GCA_902167145.1 |

In our experiments, we found a significant linear correlation between the apparent chromosome area and the DNA content. Seemingly, this stands in contrast to other results about chromosome scaling laws [23, 10, 5] which find non-linear power laws for the area. However, we analyze chromosome scaling in a single karyotype, whereas the above-mentioned studies examine chromosome scaling between different species and karyotypes. It is a classical result that chromosomes in a single metaphase spread have similar widths [13, 2, 4]. Assuming that the volume scales linearly with the DNA content, we also get a linear relationship between area and DNA content.

We proposed a simple additive error model 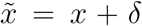 for the estimates 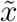 where *δ* is normal-distributed with mean *μ* = 0. This model is certainly fit for practical purposes but does not accurately describe the observed error distribution. The first observation is that the errors are, in fact, *dependent* because the estimates are always relative sizes meaning that an error in one direction in one chromosome affects all other chromosomes in the opposite direction. Secondly, we observed that a multiplicative error model 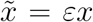 where *ε* is log-normal-distributed with expected value E[*ε*] = 1 is more accurate than the additive error model. This is equivalent to an additive normal-distributed error model in logarithmic space. Calculating the Shapiro-Wilk test statistic for normality [44, 39], we get a better fit for this model (*W* = 0.99) compared to the additive error model (*W* = 0.96). In particular, while both models do not fully describe the error distribution, i.e. *p* is close to zero, the multiplicative error model has a higher significance *p* = 3 × 10^-3^ compared to *p* =1 × 10^-7^ in the case of the additive model. Comparing panels A and B in supplementary figure 2 reveals a slightly left-skewed, i. e. right-leaning, error distribution in linear space, while it appears more symmetric in logarithmic space. However, looking at the resulting standard deviations per chromosome (supp. fig. 2C), we observe a tendency for bigger errors in smaller chromosomes, indicating a mixed additive-multiplicative error model 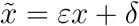. We did not investigate this model analytically because it requires non-standard statistical methods to estimate the involved distribution parameters. However, this analysis still provides valuable insights: for relatively large chromosomes, the multiplicative error term dominates the overall error, whereas the additive error term dominates the errors for relatively small chromosomes. Presumably, the multiplicative error originates from subtle difference in condensation between chromosomes and the additive error from inaccuracies of the segmentation.

## Materials and Methods

### Data Sources

KICS generally requires at least a karyotype image as input and produces estimated relative chromosome sizes as output. To acquire karyotype images, we used web searches for the species name with additional keywords like “karyotype”, “cytogenetics”, “chromosomes”, or “G-stained”. Absolute chromosome sizes can be computed using a conversion factor derived from an estimate of the genome size such as the assembly size or estimates from a genome size database like the Animal Genome Size Database [15], the Fungal Genome Size Database [25], the Plant DNA C-values Database [38], or the Bird Chromosome Database [12]. Because these databases contain literature references, they are also a good starting point for searching karyotype images.

To evaluate the performance of KICS, we compiled two datasets called Human8 and Diversity: The Human8 dataset consists of 367 chromosomes from eight karyotype images (named karyotype A-H) of human individuals found in figure 1 of [43]. These karyotype images were acquired in clinical research and contain structural defects (translocations and deletions) in eight chromosomes. We highlighted these in our results for two reasons: first, they lead to outliers in the estimates that are independent of our method, and second, they present just the type of error that is detectable by our method. We used the scaffold sizes of the CHM13 T2T v2.0 assembly ([37], accession `GCA_009914755.4`) as reference chromosome sizes because it represents the human chromosomes from telomere to telomere. The Diversity dataset covers species from amphibians, birds, fish, insects, mammals, and plants. It contains one karyotype image and one set of reference sizes for each species. Table 1 lists the species alongside the data sources. The data sources for the examples presented in figure 5 are listed in table 2. The photomicrographs of the chromosomes of *A. mexicanum* shown in [45] could not be used for our method because they are compiled from several spreads.

**Table 2:**
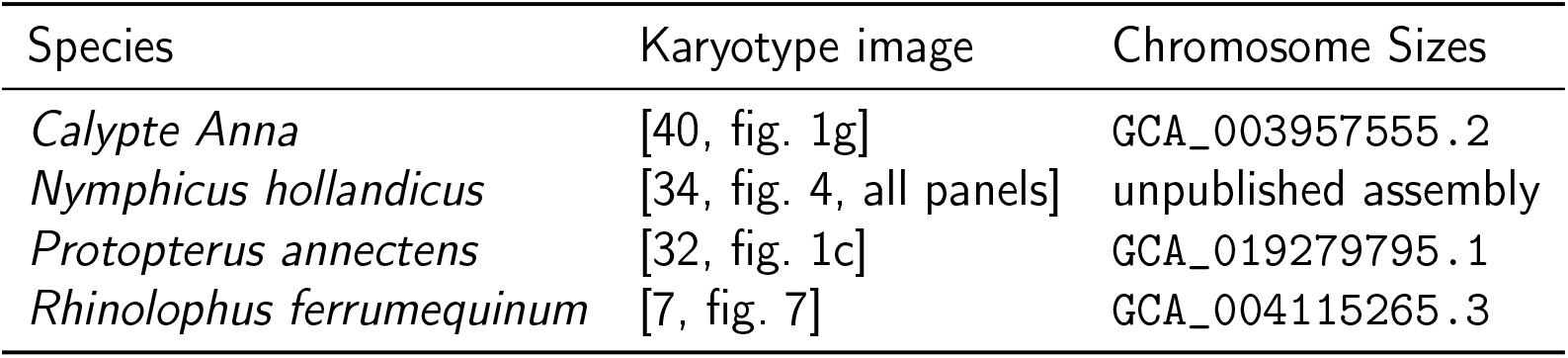
Listing of data sources for the examples. Chromosome sizes were acquired from the assemblies referenced by their GenBank accession number if given.

### Chromosome Size Estimation

#### Image Segmentation

The provided karyotype image is segmented in three phases (fig. 1.1). First, the image is converted to gray scale, where each pixel has a value in [0, 1], where 0 is the darkest and 1 the brightest value. Second, the gray-scale image is blurred with a Gaussian kernel of user-adjustable size *σ*_B_ ≥ 0. Third, the blurred image *X* = (*x_ij_*) is segmented by the threshold operation *x_ij_* < 1 – *θ*. We call *θ* the *threshold*. In our implementation, these operations are implicitly executed every time the user adjusts any of the parameters.

The user should start by selecting a threshold such that the segmentation agrees with the chromosomes. The blurring radius *σ*_B_ may be adjusted to improve the smoothness of the segmented areas, e.g., to compensate for jagged outlines. Areas that are segmented because of embedded annotations or other noise will be removed in the next step.

#### Labeling the Chromosomes

Once the basic segmentation is established, all 4-connected components are identified and labeled (fig 1.2). The background label is 0 and the foreground components are assigned labels 1, 2, 3,…. This label image is the basis for manual curation by the user. Typically, the user has to (1) assign different labels to falsely joined chromosomes (fig. 1A), (2) assign separate parts of chromosomes to the same label (fig. 1B), and (3) remove undesired labels, e. g. noise-induced segmentatmion or embedded labeling (fig. 1C). The core software `napari` provides “painting” tools (eraser, brush, fill bucket, pipette) for manipulating the label layer. Also, the user may easily remove small, noise-induced labels by selecting the n smallest labels in the table and pressing the backspace key.

#### Annotating the Chromosomes

With the image labels in place, the chromosomes can be meaningfully named (fig 1.4). Usually, chromosome names are mandatory because they carry information about the number of copies for each chromosome which is used to calculate their sizes correctly. Initial chromosomes names can be either generated automatically or by interactively striking them off in the desired order. The initial names can be manually curated using the interactive table.

The automatic labeling procedure tries to identify rows of chromosomes and in each row groups of chromosomes. The chromosome groups get numerical labels (01, 02,…) starting in the top-left corner and proceeding in rows to the bottom-right corner. The chromosomes in each group are labeled from left to right with lowercase Latin letters (a, b,…).

Annotated areas that overlap on the vertical image axis form the rows. Objects within a cutoff distance *d\** grouped together, whereas further apart objects are placed in distinct groups. The cutoff distance d* is determined for each row separately. Assume there are *n* objects in a row. Then the distances *d*_1_,…, *d*_n-1_ are given as *d_i_* = *l*_*i*+1_ – *r*_i_ where *l_i_* and *r_i_* are the left- and right-most coordinates of object *i,* respectively. Without loss of generality, assume that *d_i_* are sorted in non-descending order *d*_1_ ≤ *d*_2_ ≤– ≤ *d_n-1_*. Then the cutoff distance is given as *d*\* = max_*i*=1,…n–2_{*d_i_* | 4*d_i_* ≤ *d*_*i*+1_} unless the set is empty, in which case *d*\* = −∞, i.e., all objects are placed in separate groups. The set may be empty because either the row contains just a single object (*n* = 1) or none of the *d*_i_ satisfies the condition. The factor 4 in the above equation was determined empirically and will not work for every dataset. Also, the layout of karyotype images varies and may render this automatism useless.

In cases where the automatic naming fails, there is an interactive tool that determines the order and grouping by letting the user draw a path over each group of chromosomes. The chromosomes are then named according to the above naming scheme in chronological order and grouped if the same stroke marked them.

Finally, the names can be manually provided or altered in the interactive table. If the user enters a name in the same format as described above, it will also be interpreted in the same way. All other formats are interpreted as strings with no further meaning and chromosomes with the same names are taken to belong to the same group. This means that multiple occurrences of the same sex chromosome should get the same name, e. g. “X”.

#### Estimation of Absolute Sizes

After correctly identifying all the chromosomes, the absolute size estimates can be computed. Suppose there are *N* distinct chromosomes 1,…,*N* and each chromosome appears *c_i_* > 0 times in the karyotype. Let *A_ij_* be the area, i. e. the number of pixels, of the *j*-th annotated object (*j* = 1,…, *c_i_*) of chromosome *i* and *G* the user-provided estimate for the haploid genome size. Then we estimate the base pairs size of the annotated objects as 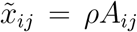, where 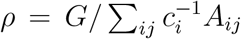 is the estimated DNA content per pixel.

Choosing the correct “genome size” may not always be straightforward. For example, the assembly in question may or may not contain both sex chromosomes, or chromosome names may be absent from a diploid karyotype impeding comparison to a haploid assembly. Generally, the “genome size” should be the number of base pairs present in the annotated image, taking into account that some chromosomes may occur multiple times. For example, a missing sex chromosome in the assembly can be compensated by skipping the annotation of that chromosome. An unannotated, diploid karyotype can be dealt with by artificially duplicating each scaffold.

### Evaluation of Experimental Results

We evaluate the accuracy and precision of KICS by matching estimated sizes to known reference chromosome sizes from independent sources. While this makes a detailed per-estimate error analysis possible, the observed errors are always a mixture of estimation errors on one side and errors in the reference sizes on the other side. Hence, we do expect to observe differences between estimates and reference sizes.

In the Human8 dataset, there are 13–16 estimates per non-defective chromosome (except for X with 10 and Y with 4 estimates). We report mean and standard deviation for each chromosome, assuming they have an additive and normal-distributed error. To assess the hypothesis of this error model, we evaluate the distribution of residuals by visually comparing it to the probability density function of a normal distribution with the parameters estimated from the samples.

Given a single karyotype image, we used the Pearson correlation as a measure of fit between the chromosome size estimates and corresponding reference sizes. We interpret values *ρ* ≤ 0.70 as insignificant, 0.75 < *ρ* ≤ 0.85 as slightly significant, 0.85 < *ρ* ≤ 0.90 as moderately significant, 0.90 < *ρ* ≤ 0.95 as very significant, and *ρ* >0.95 as highly significant. This interpretation is based on the observations we made in the course of this work.

Given estimates from a single karyotype image with unknown matching to the scaffold sizes, as typically encountered in applications of our method, we match the estimates sorted by size (largest first) with the largest scaffold sizes sorted equally but independently by size. We do not compute Pearson correlation for this data because the independent sorting introduces a strong correlation into every dataset. Instead, we evaluate the results visually by plotting the values side by side in a bar plot.

## Declarations

### Availability of Data and Materials

Karyotype images analyzed during this study are copyright protected and may be accessed through the original publications referenced in this published article. Remaining data generated or analyzed during this study are included in this published article and its supplementary information files.

### Availability and Requirements

Project name: KICS

Project home page: https://github.com/mpicbg-csbd/napari-kics

Operating systems: Platform independent

Programming language: Python

Other requirements: `napari` [9]

License: MIT

RRID: SCR_022224

biotools: napari-kics

## Abbreviations

KICS: Karyotype image-based chromosome size estimator

## Author Contributions

AL and AD implemented the software and analyzed the data. MP and GM conceived this study. MP applied and validated the software. GM supervised this study. AL wrote the manuscript. All authors read and approved the final manuscript.

## Acknowledgments

Not applicable.

