## Supplementary Figures for "Estimating chromosome sizes from karyotype images enables validation of *de novo* assemblies"

May 22, 2022
Revision 1

*Equal contribution
†To whom correspondence should be addressed:

Martin Pippel
Max Planck Institute of Molecular Cell Biology and Genetics
Pfotenhauerstr. 108
01307 Dresden, Germany
Mail:

[1]Max Planck Institute of Molecular Cell Biology and Genetics, Pfotenhauerstr. 108, 01307 Dresden, Germany
[2]Center for Systems Biology Dresden, Pfotenhauerstr. 108, 01307 Dresden, Germany

### Supplementary Figures

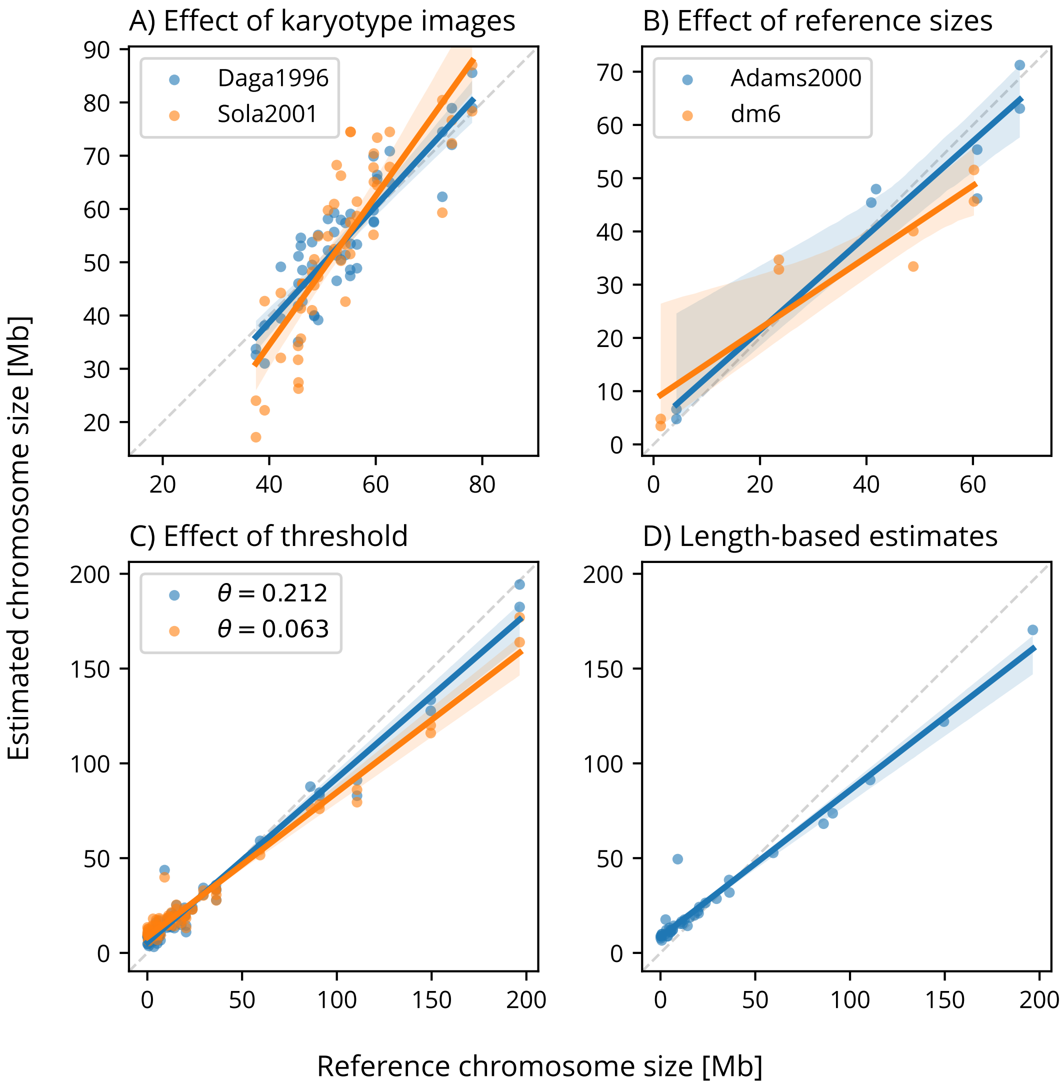


Supplementary Figure 1: Examples for the effects of varying quality of analysis inputs. **A)** Estimate results for *D. rerio* with karyotype images from Daga1996 [2] and Sola2001 [4]. **B)** Estimate results for *D. melanogaster* with reference sizes from Adams2000 [1] and the current reference assembly dm6 (`GCA_000001215.4`). **C)** Estimate results for *G. gallus* with thresholds $\theta = 0.212$ and $\theta = 0.063$. **D)** Estimates based on chromosome lengths found in [3].

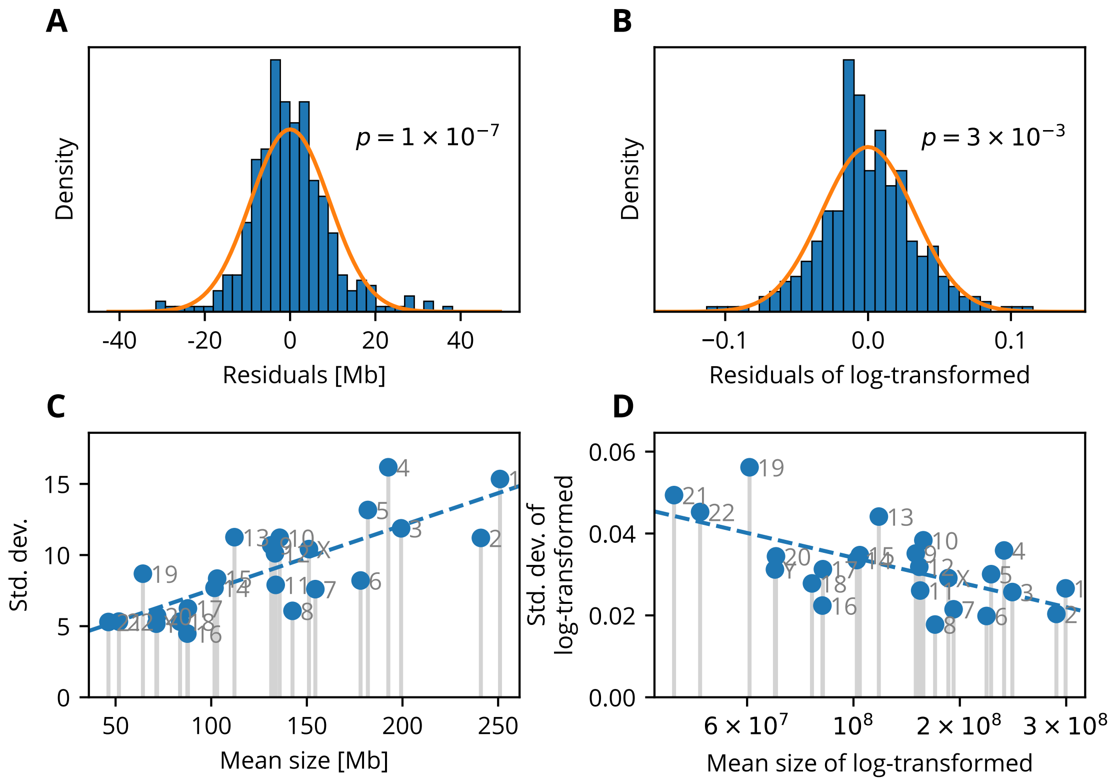


Supplementary Figure 2: Analysis of residual distribution using the Human8 dataset. The $p$-values stated in the plots are derived from the Shapiro-Wilk test. **A)** Distribution of residuals of plain chromosome size estimates. **B)** Distribution of residuals of log-transformed chromosome size estimates. **C)** Standard deviation over mean of plain chromosome size estimates. The dashed regression line highlights a trend that larger chromosomes have a higher standard deviation. **D)** Standard deviation over mean of log-transformed chromosome size estimates. The dashed regression line highlights a trend that smaller chromosomes have a higher standard deviation after log-transform.
